# Ionizing radiation has negligible effects on the age, telomere length, and corticosterone levels of Chornobyl tree frogs

**DOI:** 10.1101/2024.05.07.592866

**Authors:** Pablo Burraco, Caitlin Gabor, Amanda Bryant, Vanessa Gardette, Thierry Lengagne, Jean-Marc Bonzom, Germán Orizaola

## Abstract

Pollutants, such as ionizing radiation, released at high levels by human activities can shape ecological and evolutionary processes. The accident occurred at Chornobyl nuclear power plant (Ukraine, April 1986) contaminated a large extension of territory after the deposition of radioactive material. Beyond the immediate negative impact caused by the accident, it is still under debate whether the chronic exposure to the radiation levels currently present in the area signifies a serious threat for organisms. One hypothesis suggests that current levels of radiation may cause unobservable damage in the short-term, but have long-term effects such as decreases in longevity. Here, we investigate through a field-based approach, whether current levels of radiation in Chornobyl negatively impact the age of a semi-aquatic vertebrate, the Eastern tree frog *Hyla orientalis*. We also explore whether radiation induces changes in an ageing marker, telomere length, or in the stress hormone corticosterone. We found no effect of total individual absorbed radiation (including both external and internal exposure) on frog age (n = 197 individuals sampled in three consecutive years). We also did not find any relationship between individual absorbed radiation and telomere length, but a negative relationship between individual absorbed radiation and corticosterone levels. Our results suggest that radiation levels currently experienced by tree frogs in Chornobyl may not be high enough to cause severe chronic damage to semi-aquatic vertebrates such as this species. This is the first study addressing age and stress hormones in Chornobyl wildlife, and thus future research will confirm if these results can be extended to other taxa.

## Introduction

Human activities are altering ecosystems worldwide and, particularly, the presence of pollutants in natural environments is leading to biodiversity loss [1]. The release of high levels of ionizing radiation is a rare but potentially threatening pollutant for wildlife that can be found in certain areas of the planet as a result of human actions such as nuclear weapons tests or derived from nuclear accidents (Beresford et al. 2020). Investigating the effects of ionizing radiation on fitness components and health-related physiological parameters can provide insights into the long-term impact of radioactive contamination on biodiversity.

Ionizing radiation can directly damage DNA or interact with other molecules in the cell, causing indirect damage to DNA and even compromise organ function or individual survival [3]. Radiation can also lead to the upregulation of repair or buffering pathways such DNA repair mechanisms or antioxidants, however, these responses can be energetically costly and insufficient to fully avoid radiation effects, which could therefore result in lifespan reductions [4]. The accident at the Chornobyl nuclear power plant, on 26 April 1986, led to the largest release of radioactive material to the environment ever recorded. While the consequences of this accident were severe in the short term for both humans and wildlife (Mettler, Gus’kova, and Gusev 2007; Beresford and Copplestone 2011), it remains elusive whether current radiation levels have the potential to shape the ecology and evolution of wild populations in the contaminated areas. Recent research has reported a wide diversity of consequences of Chornobyl radiation on wild organisms, from severe impacts at the genetic and physiological level [e.g., 7–9] to lack of effects, and even the presence of abundant wildlife populations [10–13]. More than three decades have passed since the accident, and radiation levels have dropped more than 90% overall, and short-lived radionuclides known to induce significant biological damage have completely disappeared from the area (e.g. ^131^I; Intelligence Systems GEO 2011). There is a clear need for further studies aiming to address whether chronic exposure to current Chornobyl radiation (i.e., currently much lower than at the time of the accident and non-lethal in the short term) can lead to changes in animal life histories and health.

Organisms permanently exposed to ionizing radiation may experience negative effects on different fitness components such as survival and reproduction [e.g., 14]. However, no field-based study has addressed whether chronic exposure to radiation shapes age in Chornobyl wildlife, so far. Shifts in individual age can have demographic implications including effects on population abundance [15,16], and can inform about the cumulative impact of coping with radiation over the course of life. This information can be complemented with markers of individual ageing or stress such as telomere length and corticosterone levels, respectively. Telomeres are the non-coding terminal sequences of the chromosomes that shorten as a consequence of environmental harness, and thus telomere shortening is often considered a marker of organismal ageing [17]. Anthropogenic disturbances are known to have a negative impact on the telomeres of some taxa (Louzon et al. 2019; Salmón and Burraco 2022), including ionizing radiation both on wildlife and humans (Genescà et al. 2006; Kesäniemi et al. 2019; Wong et al. 2000; but see Cunningham et al. 2021). However, this process seems to be context dependent [reviewed in 19]. Likewise, corticosterone, the main glucocorticoid in amphibians, plays a key regulatory role in stress-induced responses [20,21,25]. Elevated chronic corticosterone levels can enhance metabolism and induce immunosuppression, oxidative stress, and finally have a negative impact on individual performance [26–29]. Investigating both telomere length and corticosterone levels can help to understand effects of radiation on animal ageing and health.

Here, we investigate whether current radiation levels in Chornobyl shape the age, telomere length, and corticosterone levels of the Eastern tree frog *Hyla orientalis*. Our previous results on the effects of radiocontamination on *H. orientalis* in the Chornobyl region, conducted more than 30 years after the Chornobyl nuclear accident shown no significant effects of radiocontamination on several physiological parameters. This includes unaltered blood biochemistry [12], markers of liver function [30], or the redox status of the frogs [31]. However, we detected an elevated mutation rate in the frogs living in the area compared to other European populations, with notably the presence of stop-gained mutations in the most contaminated areas and changes in the transcriptional profile in genes involved in energetic metabolism [32,33]. In this context, a chronic exposure to (relatively) low levels of ionizing radiation present in the Chornobyl region could have an impact on the lifespan of individuals. To explore this idea, we carried out an extensive sampling over three consecutive years in which we quantified individual age and absorbed dose rates in tree frogs inhabiting a gradient of radiation across the Chornobyl area. In a subset of individuals, we investigated individual ageing rate through estimates of telomere length, and recorded levels of the stress hormone corticosterone. We expected that highly radiocontaminated frogs would be, overall, younger than those experiencing lower radiation levels. We also predicted shorter telomeres and higher corticosterone levels in frogs coping with high radiation levels.

## Methods

### Field sampling

The Eastern tree frog (*Hyla orientalis*) is a cryptic species of the European tree frog (*Hyla* arborea) group found in Asia Minor and southeastern Europe [34]. We captured reproductive adult males of this species in ponds located in the Chornobyl area (both within and outside the Chornobyl Exclusion Zone; see Figure 1) in mid-May across three consecutive years (i.e. 2016-2018). We sampled 84 frogs in 2016, 70 in 2017, and 102 in 2018, for a total of 256 individuals sampled in 14 different localities across a marked gradient of ambient radiation within the Chornobyl area (see Figure 1 and Table S1). Total individual absorbed dose rates, including external and internal exposure levels (and both ^137^Cs and ^90^Sr) were estimated for all frogs (see Burraco, Car, et al. 2021 for details). Although calculating total individual absorbed dose rates is the most accurate method to estimate radiation levels to which an individual has been exposed throughout its lifetime (Beresford et al. 2020), we also used ambient radiation levels to classify the localities included in this study in two categories: low (0.01-0.27 µSv/h) and medium-high radiation (1.09-32.4 µSv/h; see Burraco, Car et al. 2021 for methodological details). We used this second approach to compare the results in our study system with previous ecological studies conducted in the area (e.g. Møller and Mousseau 2006; 2016; Beresford et al. 2016).

**Figure 1.**
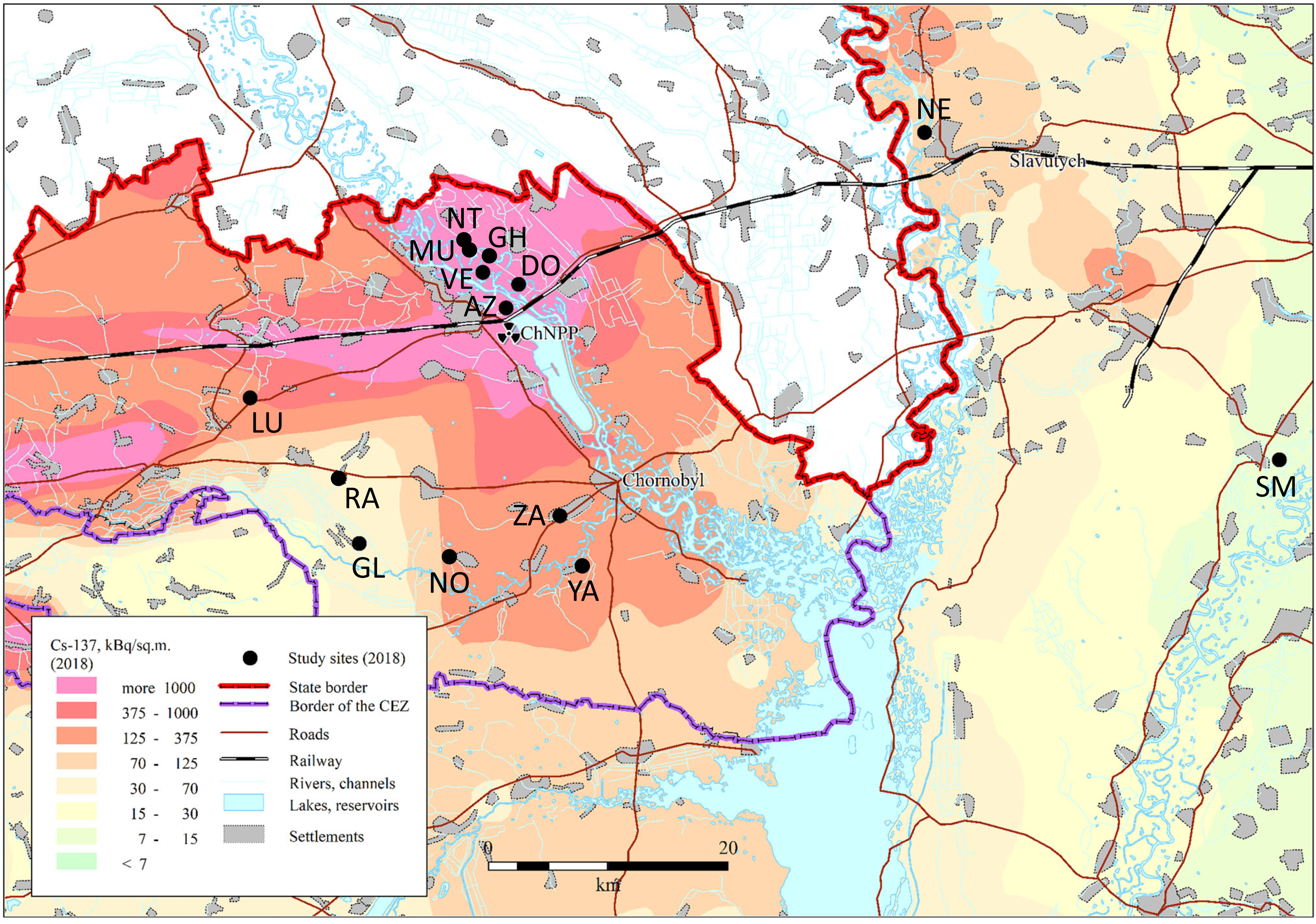
Map of the studied Eastern tree frog (*Hyla orientalis*) locations (see also Table S1). The underlying ^137^Cs soil data (decay corrected to spring 2018) is derived from the atlas of radioactive contamination of Ukraine (Intelligence Systems GEO, 2011).

Frogs were captured during the night, transported to the laboratory located in Chornobyl town (Ukraine), and individually placed in buckets containing 5 cm of clean water until the next morning. Then, we recorded individual snout-to-vent length and width using a caliper, and we weighed each individual using a precision balance to the nearest 0.01 g. We euthanized frogs by pithing without decapitation [following AVMA guidelines: ,39]. From each frog, we collected the femur of the right hind limb for age estimation, and stored it in ethanol 70% until processing. In 2018, before euthanasia, we introduced a small ball of cotton in the mouth of frogs for ∼15 secs for saliva corticosterone quantification (N = 54 frogs). We also collected a portion of muscle from the right hind limb for telomere length quantification (N = 62 frogs). All animals were collected and experimental procedures conducted, under permit of Ministry of Ecology and Natural Resources of Ukraine (No. 517, 21.04.2016).

### Individual age

Skeletochronology allows estimating the age of animals with indeterminate growth living in temperate climates, by making use of the annual growth rings formed when osteogenesis is low or inactive. Such rings are defined by lines called *lines of arrested growth* (LAGs). Males of *H. orientalis* are mostly inactive during the cold months, which makes them good candidates for skeletochronolgy studies [40,valited for Hyla frogs by 41].

We conducted age estimations at the Laboratorie d’Écologie des Hydrosystèmes Naturels et Anthropisés (Université Claude Bernard – Lyon I, CNRS, France). We sampled femur bones that were decalcified in 1% nitric acid for 20 minutes and rinsed. Cross sections of the diaphyseal region of the bone were obtained using a freezing microtome (Microtom heidelberg HM330). We obtained several 18-20-micron sections and stained them with Ehrlich’s haematoxylin. We placed them onto microscope slides. We examined sections and quantified the number of LAGs on one section using a light microscope (Olympus CX40). Individual age was estimated by counting the number of lines following the first identifiable LAG (always by the same researcher, VG), which is preceded by a well-developed layer of bone [42]. The LAG of the first winter after metamorphosis is assumed to be very thin and is likely to be reabsorbed [42], therefore we added one year to the counted number of LAG.

### Relative telomere length

We conducted telomere analyses at the physiology laboratory of the Doñana Biological Station (Spanish National Research Council, CSIC). We isolated genomic DNA using a commercial kit (QIAGEN DNAeasy Blood&Tissue Kit) and stored it at -80 °C until assayed. We estimated relative telomere length through real-time quantitative PCR. This method compares the amount of telomeric sequences relative to a control single-copy gene [43,44]. In our study, we amplified of the ribosomal 18S gene as the qPCR control gene, using the following primer combination: forward 5-ACTCAACACGGGAAACCTCA-3′ and reverse 5′-AACCAGACAAATCGCTCCAC-3′. To amplify the telomeric region, we used primers designed for vertebrates: tel1b 5′CGGTTTGTTTGGGTTTGGGTTTGGGTTTGGGTTTGGGTT-3′ and tel2b 5′-GGCTTGCCTTACCCTTACCCTTACCCTTACCCTTACCCT-3′. Each sequence was amplified in a separate qPCR plate using Bio-Rad CFX96 Real Time thermal cycler. For 18S amplification, we prepared a master mix containing 10 μL of SYBR Green QPCR Master Mix (Applied Biosystems), 1.2 μL of each primer for a final concentration of 300nM, 1 μL of genomic DNA at 5 ng/μL, and 6.6 μL of water. For the telomere sequence, we prepared a master mix consisting in 10 μL of SYBR Green QPCR Master Mix (Applied Biosystems), 0.4 μL of each primer for a final concentration of 100nM, 1 μL of genomic DNA at 5 ng/μL, and 8.2 μL of water. All qPCRs started with a temperature profile of 10 min at 95°C followed by 40 cycles of 10 seconds at 95°C, 15 seconds at 65°C, and 15 seconds at 72°C for both genes. Each plate included a standard curve with five points of a serially diluted from a sample containing a pool of samples from different sampling sites and years. The determination of Cycle threshold (C_t_) values was performed with the CFX Manager Software (Bio-Rad). We calculated plate efficiencies and ran melt curves demonstrating single peaks. We ran all samples by duplicate and calculated relative telomere length according to [45]. The intra-plate CV% was, respectively, 0.87% and 1.56% for 18S and telomere amplifications. Efficiency was, respectively, 98.11% and 94.92% for 18S and telomere amplifications. For each gene, samples were run in a single 384-well plate.

### Corticosterone

Saliva samples were shipped to the Department of Biology, Texas State University (USA) for corticosterone determination. Here, we extracted and quantified saliva CORT following [46]. To quantify saliva volume, individual swabs were thawed, placed above a plastic filter in microcentrifuge tubes, and centrifuged at 10,000 rpm for 10 min. We measured saliva volume from the bottom of each tube using a syringe (rinsed three times with EtOH and then three times with ddH2O between samples) and left it in the original microcentrifuge tube. Mean saliva volume was 3.27 ± 0.44 μl. We then washed each swab with 300 μl of EIA buffer into the original microcentrifuge tube, to increase yield of CORT from saliva swabs and centrifuged again at 10,000 rpm for 10 min. We added trichloroacetic acid (TCA) to precipitate salivary proteins at a ratio of 20% of saliva volume (Hammond et al., 2020). We then vortexed samples for 10 s, incubated at room temperature for 30 min, centrifuged at 10,000 rpm for 10 min, and finally supernatants were transferred to a new microcentrifuge tube. We stored purified and concentrated samples at −20 °C until assayed. We used a commercial EIA kit to quantify salivary CORT (item no. 501320, Cayman Chemical Company). Samples were assayed in duplicate read on a spectrophotometer plate reader at 405 nm (BioTek 800XS). Inter plate variation was 8.82% for two plates. Methods to quantify corticosterone concentration in saliva have been validated in several amphibian species [46–48].

### Statistical analyses

All analyses were conducted in R software (version 4.2.1). We checked for parametric assumptions through Kolmogorov-Smirnov and Breusch-Pagan tests. We obtained individual body condition estimates by getting the residuals of the linear relationship between body mass and length [49]. Since body condition is known to correlates with absorbed dose rates in Chornobyl tree frogs [33], we included this parameter in the analyses. To study the relationship between individual absorbed dose rate (i.e., as the predictor variable) and age, we conducted a generalized model with a Poisson distribution, including as fixed factors GPS coordinates where samples were collected to correct by spatial correlation, body condition, and year of sampling. To investigate the effect of ambient radiation on age, we conducted a generalized model with Poisson distribution, including age as the dependent variable, ambient radiation category as the independent category, and GPS coordinates, body condition, and year of sampling as fixed factors. We tested for the relationships between telomere length, individual absorbed dose rate, and age through a linear model including the three variables, and GPS coordinates and body condition as covariates. Finally, we ran a linear model to test for an effect of individual absorbed dose rate on corticosterone levels, including GPS coordinates, age, and body condition (except for corticosterone because values were adjusted to individual body mass) as covariates. We only included age estimates considered to have a high degree of reliability (197 out of 256 sampled individuals).

## Results

Frog age varied between 2 and 9 years (Table S1; Figure 2). Body condition and age showed a significant positive relationship (*Χ_1,177_* = 5.82, *P*-value = 0.016). We found no significant correlation between total individual absorbed dose rate and frog age (*Χ_1,177_* = 2.27, *P*-value = 0.132, R-squared = 0.05; Figure 2A). Frogs inhabiting locations with medium-high ambient radiation levels were significantly younger (4.44 vs 4.82 years, i.e., 9.95% younger on average; Figure 2B) than those from populations with levels close to background (‘radiation category’ effect: *Χ_1,177_* = 6.71, *P*-value = 0.009, Figure 2B). Relative telomere length was not affected by individual absorbed dose rates (*F_1,57_* = 1.69, *P*-value = 0.194; Figure 3A), nor age (*F_1,57_* = 0.13, *P*-value = 0.572). Finally, corticosterone levels were negatively affected by individual absorbed dose rates (*F_1,51_* = 4.22, *P*-value = 0.044; Figure 3b), and it was not affected by age (*F_1,51_* = 0.01, *P*-value = 0.988).

**Figure 2.**
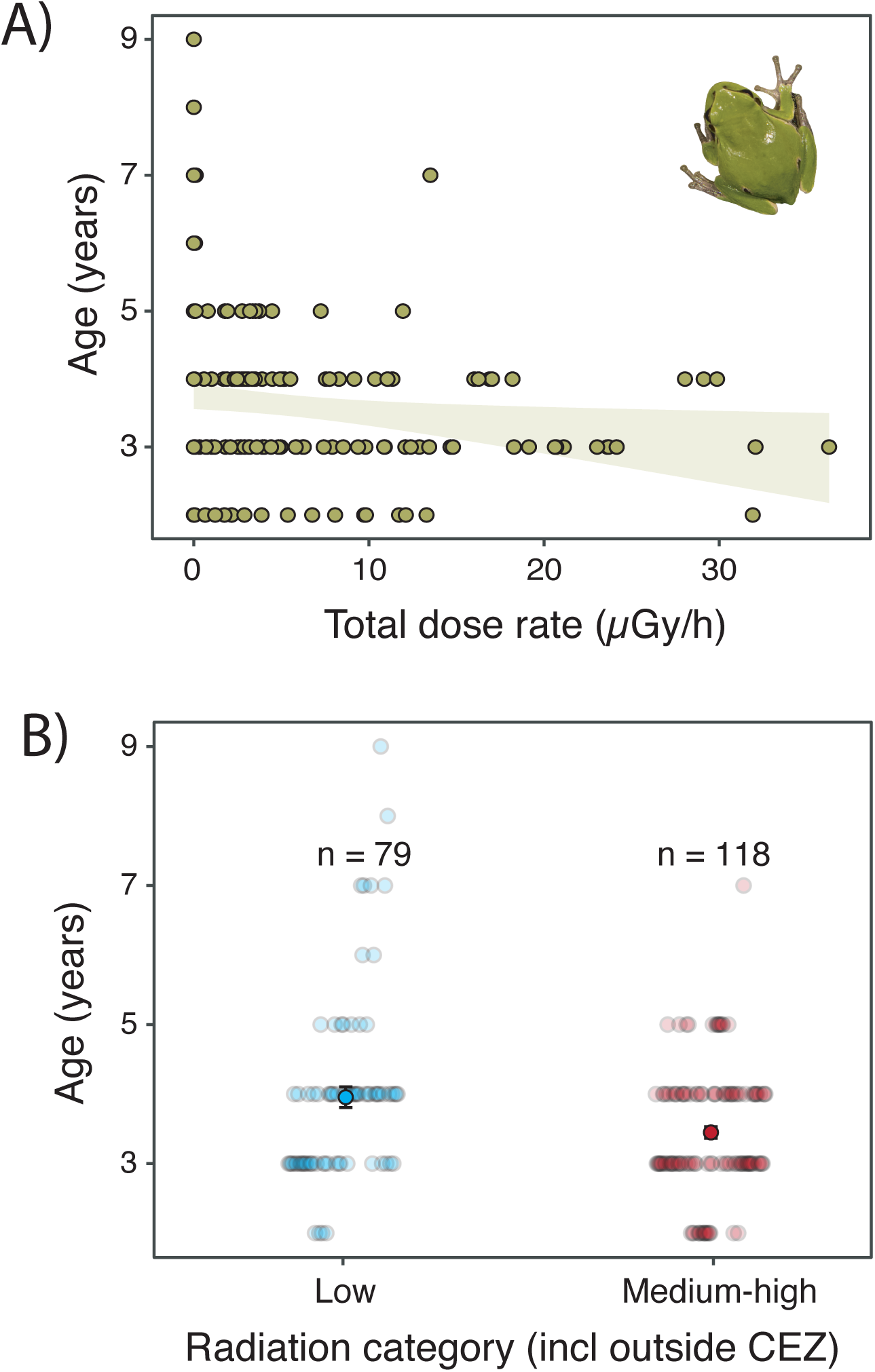
A) Correlation between frog age and total individual dose rates in Eastern tree frog (*Hyla orientalis*) males sampled within the Chornobyl area (see map in Figure 1). Since the correlation is not significant, regression line is not drawn, grey lines indicate the 95% confidence interval. B) Age difference between frogs inhabiting localities with low (0.01-0.27 µSv/h) and medium-high radiation (1.09-32.4 µSv/h).

**Figure 3.**
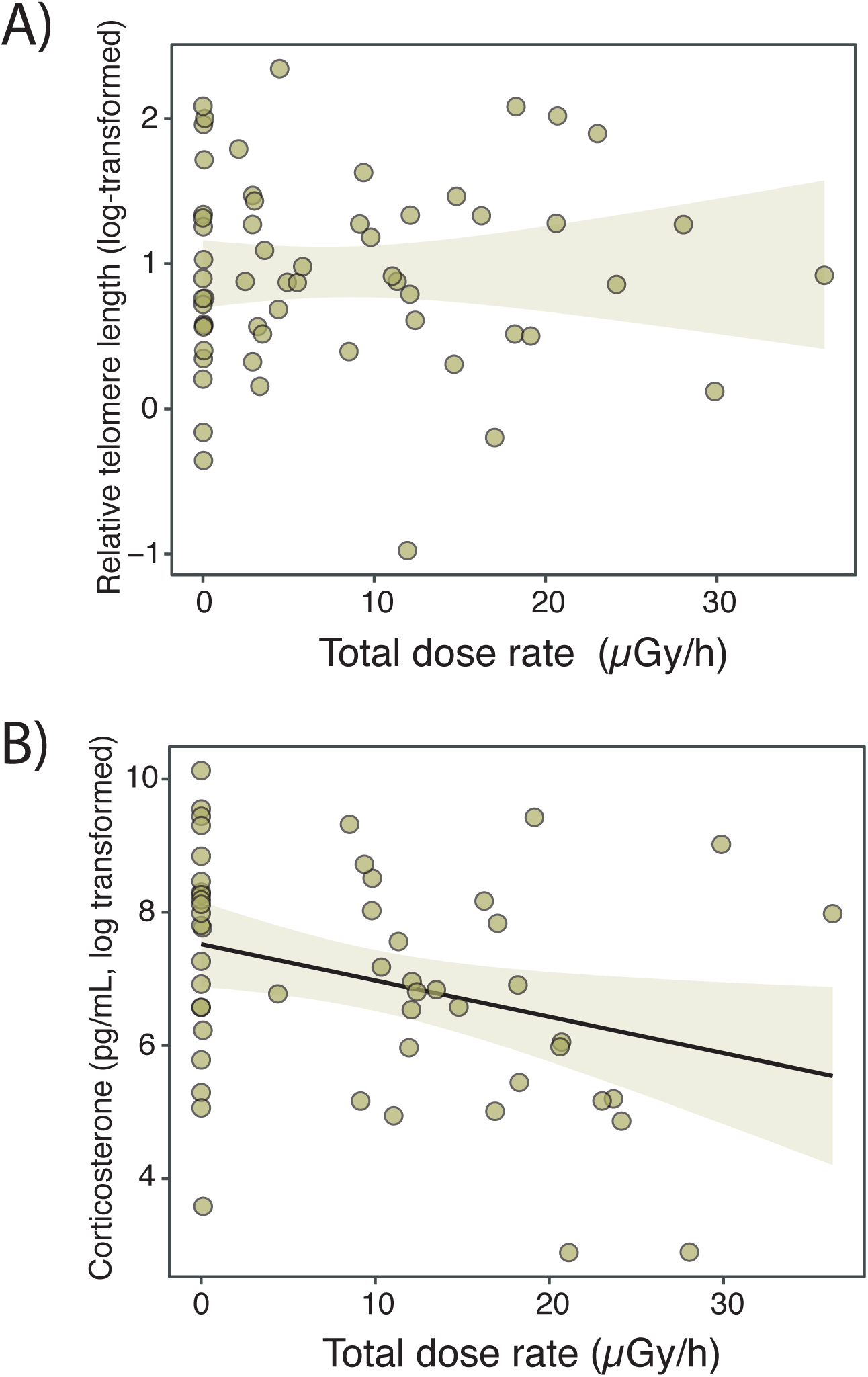
Correlations between frog age and (A) relative telomere length, and (B) corticosterone levels in Eastern tree frog (*Hyla orientalis*) males sampled within the Chornobyl area (see map in Figure 1). Regression line is drawn only when the correlation is significant, grey lines indicate the 95% confidence intervals.

## Discussion

Our study shows that radiation levels absorbed by tree frogs in Chornobyl had no effect on the age of the frogs. When considering only ambient radiation, individuals from localities with high radiation levels were, overall, slightly younger than frogs from localities much less affected by radiation. Radiation did not affect ageing rates in relation to the lack of variation in telomere length among individuals. Frogs experiencing high dose rates had lower levels of the stress hormone corticosterone. This research represents the first approach for the study of the effects of chronic radiation exposure on the age and stress hormones in Chornobyl wildlife.

The magnitude of the impact caused by the chronic exposure to radiation in an organism is driven by the total dose absorbed by an organism over a period of time. In Chornobyl, radiation has decreased considerably from the moment of the accident [50], and nowadays there is controversy about the effects of current radiation levels on wildlife [e.g., 2]. In our study system, Chornobyl tree frogs inhabiting radiocontaminated areas show normal physiological condition [12,30,31,but see 32,33 addressing effects at the genetic or transcriptomic level]. In accordance with our previous physiological approaches in the same study system, we did not find a marked relationship between individual absorbed dose rates and frog age. Furthermore, our findings are based on estimates of the radiation absorbed by each individual frog, including both external and internal radiation [for more details, see 35]. Our study suggests that current radiation levels experienced by tree frogs in Chornobyl are not enough to markedly shorten their lifespan, and agree with a previous study conducted on a similar species in radiocontaminated areas around Fukushima [Japan, 51]. Also, previous studies have shown that radiation doses currently experienced by tree frogs in Chornobyl are below the threshold considered harmful for amphibians [35], which may explain the little impact observed on the age of frogs.

When considering only environmental radiation levels, as done in many other studies in Chornobyl [36–38], and grouping localities with contrasting radiation levels (i.e. low and medium-high radiation), we found that frogs were slightly younger in highly-contaminated localities than in those areas with radiation levels currently close to background. This result might suggest negative organismal impacts of early-life environmental conditions, parental effect or selective disappearance. However, the use of environmental radiation prevents from considering differences in individual exposure to radiation. Nevertheless, the overall small effect of radiation on frog age may suggest negligible ecologically-relevant effects on this trait. Age values recorded in our study are within the age range observed in the same species or in closely-related species such as *Hyla arborea* in areas never exposed to radioactive fallout [41,52–54]. Hence, it seems that radiation has not caused a decrease in this crucial life-history trait, more than 30 years after the accident. A study conducted early after the accident showed a marked reduction in lifespan in experimental white mongrel rats from Chornobyl town and exposed to high radiation levels [55]. In humans, studies have shown, for example, evidence of increased risk of hematological malignancies, and the development of cataracts or cardiovascular diseases caused by radiation [56,57], but the effect of chronic radiation exposure on the age structure of human populations has not been investigated yet. Long-term research should focus on increasing our understanding of possible age-dependent effects of chronic exposure to radiation on the fitness and health of different taxa [58].

Individual absorbed dose rates also do not seem to impact on the ageing rate of Chornobyl tree frogs, based on its relation to telomere length. Since radiation does not seem to affect frog age, the lack of correlation between radiation and telomere length may further support that finding. The actions of different mechanisms protecting from telomere attrition such as the enzyme telomerase or the antioxidant machinery, might be also behind the observed pattern. Telomerase expression is upregulated in some tissues in bank voles coping with Chornobyl radiation [23]. Likewise, Chornobyl tree frogs with high dose rates do not experience higher lipid peroxidation levels [31], which may indicate physiological compensatory responses to radiation, similar to those observed in other amphibians coping with detrimental conditions earlier in their life [59]. We should also notice that we measured telomere length in the hindlimb muscle of frogs, but telomere dynamics can be tissue-specific in amphibians and other vertebrates [23,60,61]. Indeed, telomere dynamics differ across tissues in bank voles from the Chornobyl Exclusion Zone [23]. In agreement with our results, no damage on DNA, including telomeres, was recorded in wild boars and snakes exposed to radiation released after the Fukushima nuclear accident (Japan; [62]. Finally, long term exposure to environmental perturbations may lead to chronic stress, which can result in either up or downregulated corticosterone release, both known to cause detrimental effects on organisms [63–65]. Here, we found lower chronic corticosterone levels in frogs with high absorbed radiation rates. In our study, frogs were maintained in the lab overnight before taking saliva samples for measuring the hormone, which may have induced corticosterone responses. Capture and transportation to the lab is stressful to amphibians and thus frogs from all sites may have been similarly stressed [66–68]. Hence, frogs chronically challenged with high absorbed rates may have unable to upregulate their CORT levels in response to capture. A higher release of CORT under stressful events (such as capture) has been suggested as an adaptive response to allocate energy that facilitates coping with the challenging condition [69].The study of stress hormones has been overlooked in Chornobyl research (already highlighted by Møller and Mousseau 2006), and thus future work should determine if our findings can be extended to other taxa.

## Conclusions

Our study finds no effect of ionizing radiation levels currently experienced by tree frogs inhabiting Chornobyl on their age, nor on an ageing marker, telomere length. We observed a negative relationship between individual absorbed dose rates and corticosterone levels. These results seem to be in the line of our previous work on different phenotypic traits in the same study system, and suggest that current Chornobyl radiation levels are not high enough to cause chronic damage to a semi-aquatic vertebrate such as tree frogs.

## Acknowledgements

Sergey Gashchack, Yevgenii Gulyaichenko and the staff of the Chornobyl Center for Nuclear Safety, Radioactive Waste and Radioecology (Slavutych, Ukraine) helped us with field work and radioisotope analyses. Brian Tornabene provided help with saliva corticosterone extraction methods. This work was supported by projects from the Swedish Radiation Protection Agency-SSM (SSM2018–2038) and Carl Tryggers Foundation (CT 16:344) to GO, Uppsala University Zoological Foundation, Helga Ax:son Johnsons Stiftelse, and Spanish Association of Terrestrial Ecology-AEET to PB, and the French Institute for Radiological Protection and Nuclear Safety-IRSN to JMB. PB was supported by a Carl Tryggers Foundation scholarship (CT 16:344) and by a Juan de la Cierva Incorporación funded by the Spanish Ministry of Science and Innovation (IJC2020-044682-I). JMB was financially supported by IRSN, and GO was supported by the Spanish Ministry of Science, Innovation and Universities (Ramón y Cajal Program, RYC-2016-20656).

## Data availability

A data table with all original measurements are deposited in Figshare repository 10.6084/m9.figshare.25700271

## Supplementary Material

**Table S1.** Geographic coordinates (latitude and longitude), number of frogs used in the study, current levels of environmental radiation (i.e. ambient dose rate, different among years for some localities), and tree frog age range for the locations included in the study.

| Location | Code | GPS coordinates | Sampled frogs (n) | Ambient Dose Rate ($\mu\text{Sv/h}$) | Age range (years) |
| --- | --- | --- | --- | --- | --- |
| Azbuchin | AZ | 51.4047, 30.1044 | 35 | 32.40-7.61 | 2-5 |
| Vershina | VE | 51.4328, 30.0769 | 13 | 16.20 | 3-7 |
| Muravka | MU | 51.4515, 30.0528 | 11 | 3.70 | 3-5 |
| Glyboke Hydro | GH | 51.4447, 30.0711 | 6 | 3.70 | 3-5 |
| Northern Trace | NT | 51.4567, 30.0486 | 11 | 2.51 | 3-5 |
| Dolzhikovo | DO | 51.4256, 30.1161 | 40 | 2.10-1.09 | 2-5 |
| Lubianka | LU | 51.3388, 29.7976 | 3 | 0.27 | 5-7 |
| Novosiolki | NO | 51.2195, 30.0430 | 7 | 0.13 | 3-4 |
| Zalesie | ZA | 51.2506, 30.1667 | 11 | 0.12 | 3-5 |
| Yampol | YA | 51.2119, 20.1899 | 12 | 0.10 | 3-4 |
| Glinka | GL | 51.2300, 29.9250 | 22 | 0.10 | 2-7 |
| Nedanchichy N | NE | 51.5364, 30.5981 | 6 | 0.08 | 3-4 |
| Razjezzheie | RA | 51.2786, 29.9050 | 7 | 0.07 | 3-5 |
| Smolin | SM | 51.2925, 31.0294 | 7 | 0.04 | 3-9 |

